# *De novo* germline mutation in the Dual Specificity Phosphatase 10 gene accelerates autoimmune diabetes in Non-Obese Diabetic (NOD) mice

**DOI:** 10.1101/2020.12.30.424747

**Authors:** Anne-Perrine Foray, Sophie Candon, Sara Hildebrand, Cindy Marquet, Fabrice Valette, Coralie Pecquet, Sebastien Lemoine, Francina Langa-Vives, Michael Dumas, Peipei Hu, Père Santamaria, Stephen Lyon, Lindsay Scott, Chun Hui Bu, Tao Wang, Darui Xu, Eva Marie Y. Moresco, Claudio Scazzocchio, Jean-François Bach, Bruce Beutler, Lucienne Chatenoud

**Affiliations:** Université de Paris, Paris, France; Institut Necker-Enfants Malades, CNRS UMR8253, Inserm UMR1151, Paris, France; Center for the Genetics of Host Defense, University of Texas Southwestern Medical Center, Dallas, TX 75390, USA; Mouse Genetics Engineering Center, Institut Pasteur, 75724, Paris, France; CNRS UMR7242, Biotechnology and Cell Signaling, University of Strasbourg, 67412, Illkirch, Cedex, France; Julia McFarlane Diabetes Research Centre (JMDRC) and Department of Microbiology, Immunology and Infectious Diseases, Snyder Institute for Chronic Diseases and Hotchkiss Brain Institute, Cumming School of Medicine, University of Calgary, Alberta, Canada; Quantitative Biomedical Research Center, Department of Population and Data Sciences, University of Texas Southwestern Medical Center, Dallas, TX 75390, USA; Department of Microbiology, Imperial College London, London SW7 2AZ, United Kingdom

**Author notes:** Equal contributions (Anne-Perrine Foray, Sophie Candon). Anne-Perrine Foray, Weill Cornell Medicine, New York, NY, USA, Sophie Candon, Laboratoire d’immunologie et biothérapies, CHU de Rouen, 76000, Rouen, France. Correspondence: Jean-François Bach,; Bruce Beutler,; Lucienne Chatenoud,.

## Abstract

Here we report the isolation by selective breeding of two sublines of Non-Obese Diabetic (NOD) mice exhibiting a significant difference in the incidence of autoimmune type 1 diabetes (T1D). Whole genome sequencing of the NOD/Nck^H^ (high T1D incidence) and NOD/Nck^L^ (low T1D incidence) revealed the presence of a limited number of variants specific to each subline. Treating the age of T1D onset as a quantitative trait and using automated meiotic mapping (AMM), enhanced susceptibility in the NOD/Nck^H^ subline was unambiguously attributed to a recessive allele of *Dusp10* which encodes a dual specificity phosphatase. The causative effect of the mutation was verified with a high level of confidence by targeting *Dusp10* with CRISPR/Cas9 in NOD/Nck^L^ mice: in these animals a higher incidence of diabetes was observed. Expression of wild-type *Dusp10* correlated with higher levels of surface PD-L1 in the islets of NOD/Nck^L^ mice.

---

Type one diabetes (T1D) is an autoimmune disease caused by T lymphocytes reactive with the insulin-producing β-cells of the islets of Langerhans (*1*). The etiology is only partially understood though compelling data point to the involvement of both genetic and environmental factors (*2–4*). The disease occurs spontaneously in Non-Obese Diabetic (NOD) mice. The inbred strain NOD/Shi was first established in Japan in 1980 (*5*), and from it different colonies were established over the world, including our own in Paris, hosted at the Hôpital Necker since 1986 (NOD/Nck). Rupture of self-tolerance in NOD mice is evident at 3 weeks of age. Progressive infiltration of the islets of Langerhans by mononuclear cells (i.e. insulitis) begins at this time, eventually causing the selective destruction of β-cells that precedes the advent of hyperglycemia, by 3 months of age. By this time approximately 70% of the insulin-secreting β-cell mass has been destroyed (*1*). The incidence of T1D, as assessed by the advent of glycosuria and hyperglycemia, varies among the different colonies of NOD mice a feature considered to be largely but not exclusively due to differences in the breeding environment and conditions (*3, 6, 7*).

In this study, we took advantage of a restart of our NOD/Nck colony from a single pair of mice. In breeding the colony, we noted as we had in the past that disease incidence widely differed between litters; an observation easy to monitor because the number of pups in NOD litters often ranges from 8-12. The difference was more apparent in male mice whose disease incidence rarely exceeds 50% at 40 weeks of age than in females that show a higher disease incidence (80-95%). We decided to explore whether this phenotypic variation could be maintained in sublines of NOD/Nck mice generated through brother-sister breeding from litters presenting high or low T1D incidence. We initially established 5 distinct sublines which we followed for 3 consecutive generations (Fig. 1a-b, Supp Fig 1a). We subsequently concentrated on two of these sublines showing the most significant difference in T1D incidence. These we named NOD/Nck^H^ (for High incidence) and NOD/Nck^L^ (for Low incidence) **(Figure 1)**. The phenotypic variation was stably maintained for over 30 generations in these 2 sublines raised in a specific pathogen free (SPF) environment, as assessed by regularly comparing T1D incidence and age of T1D onset at equivalent generations **(Figure 2a)**. Furthermore, we confirmed that the difference in T1D incidence in a group of NOD/Nck^H^ and NOD/Nck^L^ mice re-derived from embryos frozen at generation 7 was comparable to that of the original G7 mice **(Figure 2b)**.

**Figure 1:**
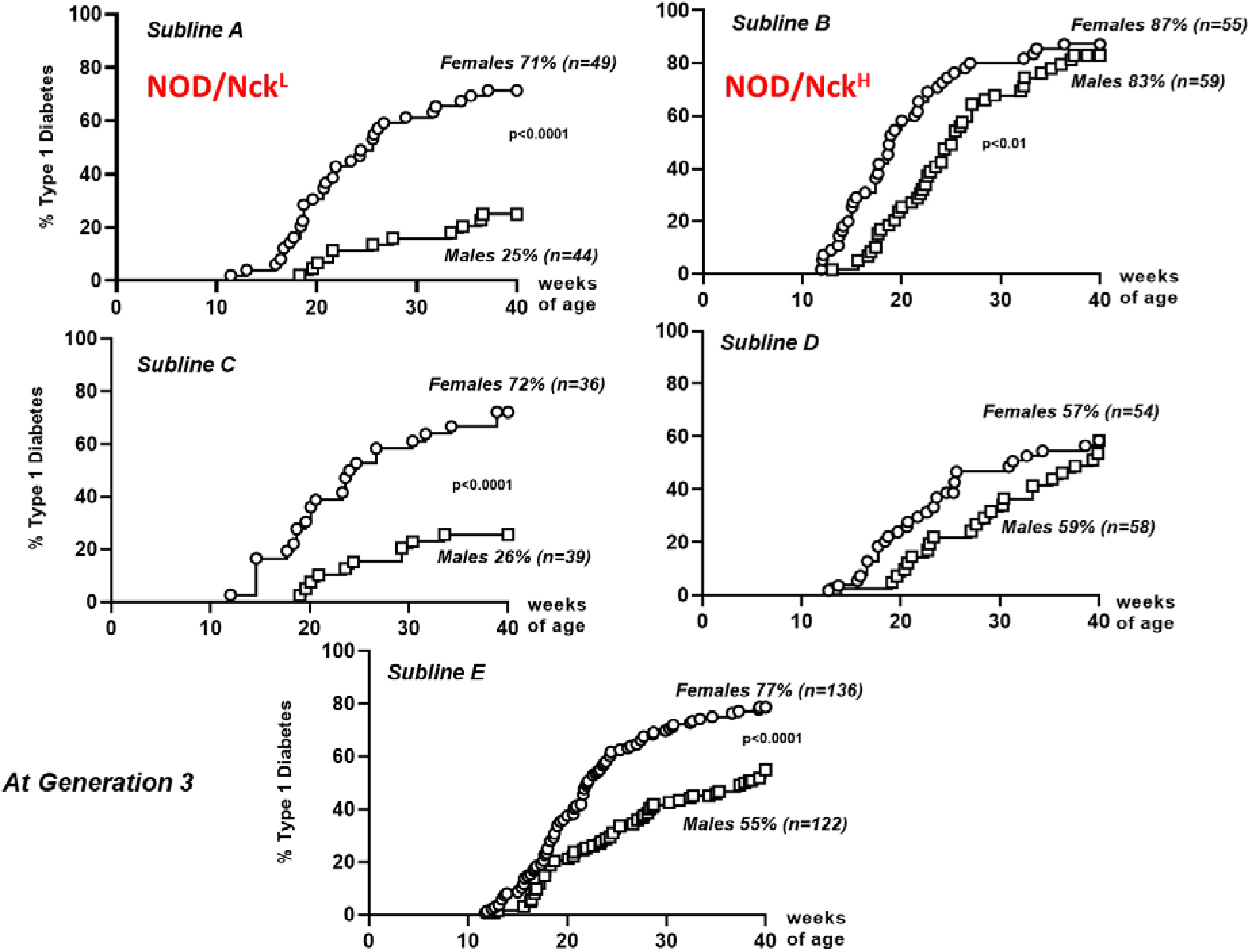
Sublines of the NOD/Nck strain. Five distinct sublines (A to E) were established from the NOD/Nck strain by brother-sister mating of mice within individual litters expressing variable T1D incidence. The five sublines were followed for three consecutive generations. For further studies we concentrated on two of these sublines: Sublines A and B which we named NOD/Nck^L^ (for low incidence) and NOD/Nck^H^ (for high incidence).

**Figure 2:**
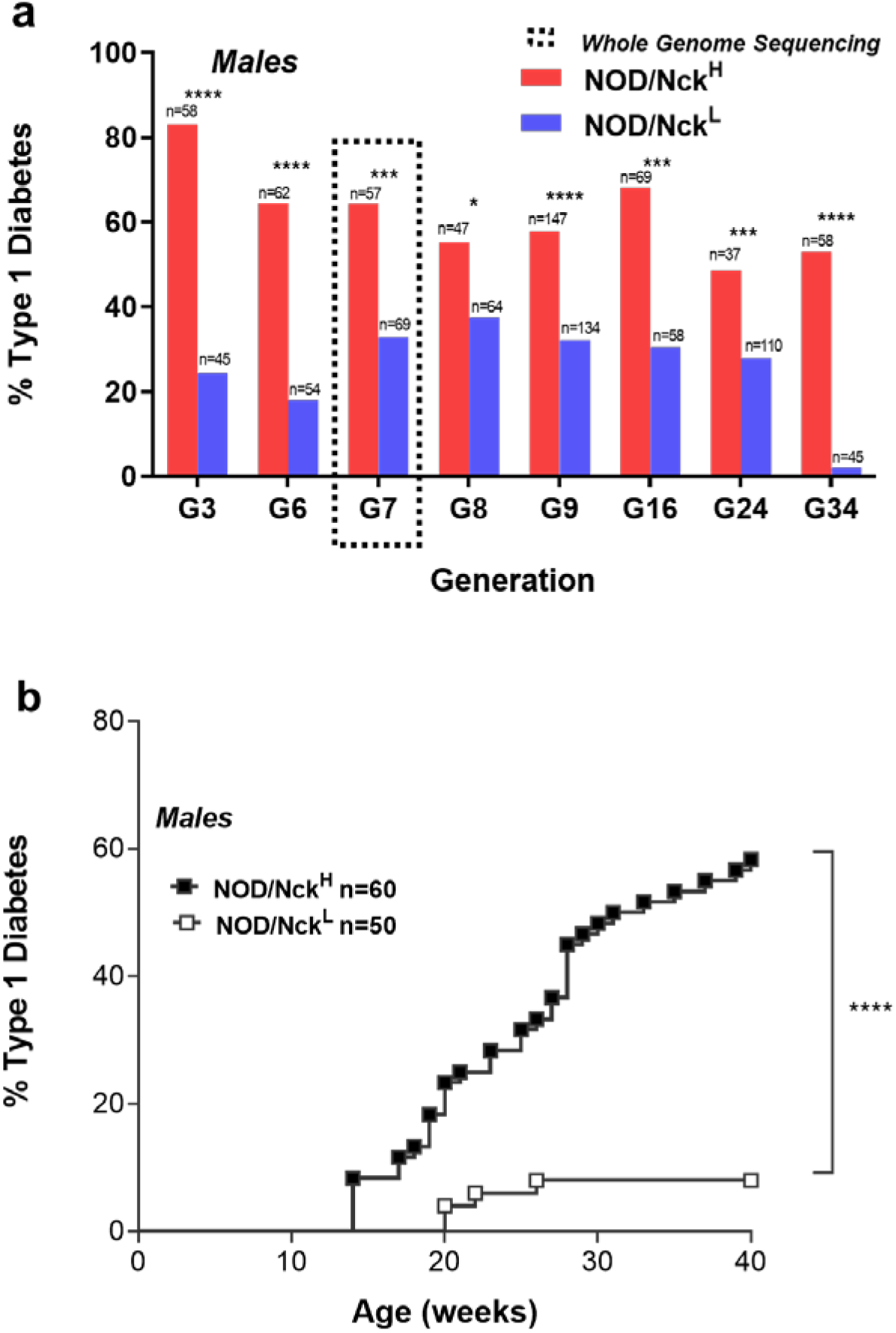
Incidence of T1D in NOD/Nck^H^ and NOD/Nck^L^ sublines. **a)** Incidence of T1D in NOD/Nck^H^ and NOD/Nck^L^ male mice followed from generation 3 up to generation 34. **b)** Incidence of T1D in NOD/Nck^H^ and NOD/Nck^L^ male mice derived from embryos frozen at generation 7 and revitalized.

Longitudinal monitoring of various immunopathological parameters characteristic of the autoimmune process in the NOD mouse showed a clear time-shifted difference in the kinetics of events that occurred earlier in NOD/Nck^H^ as compared to NOD/Nck^L^ mice. This was the case for the progression of insulitis **(Figure 3a)** and also the enumeration of diabetogenic lymphocytes in spleen, pancreatic lymph nodes and peripheral blood assessed by interferon gamma (IFNγ) ELIspot and tetramer staining respectively **(Figure 3b,c,d)**. Of note, the class I MHC tetramer complexed with the NRP mimotope (NRP-V7) that detects circulating pathogenic CD8+ T cells specific for IGRP confirmed its value as a sensitive marker to predict T1D advent in individual mice **(Figure 3e)** (*8*). However, no intrinsic abnormality in the *in vivo* functional capacity of diabetogenic T lymphocytes or regulatory CD4+FoxP3+ T lymphocytes was observed using adoptive transfer (data not shown). Similar timing of islet antigen presentation was observed in NOD/Nck^H^ and NOD/Nck^L^ mice as assessed by a well-established method: an identical capacity of diabetogenic cells expressing the transgenic BDC2.5 T cell receptor (TCR; specific for an insulin-chromogranin A fusion peptide) to migrate and proliferate in pancreatic lymph-nodes when adoptively transferred into 4-week-old recipients **(Figure 4).**

**Figure 3:**
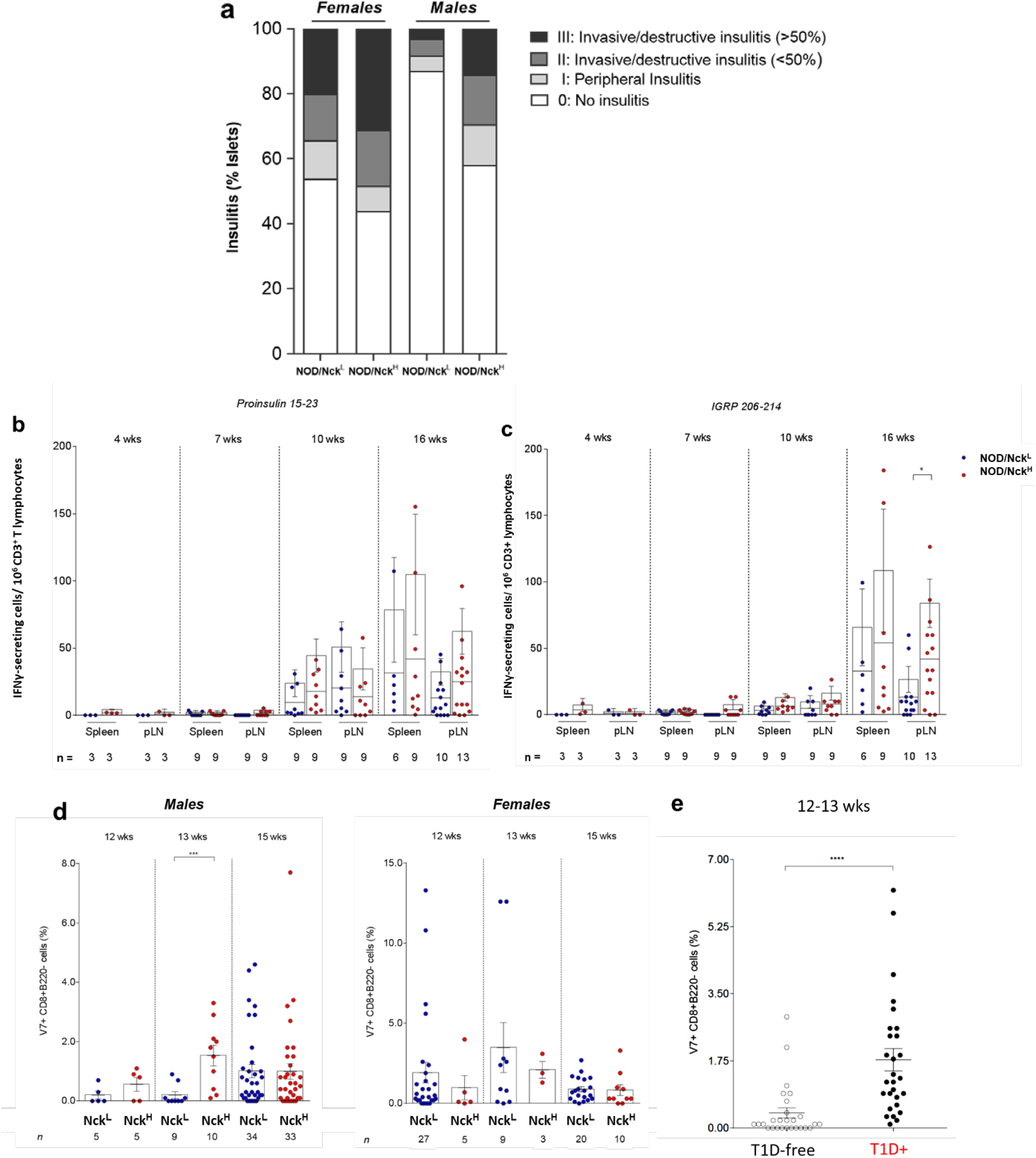
Immunological parameters characteristic of the autoimmune process in NOD mice. **a)** Insulitis in NOD/Nck^L^ and NOD/Nck^H^ mice at 12 weeks of age (n=8 in each group analyzed; a total of 60 to 80 islets were scored per mouse). The scoring system used is indicated in the legend. **b)** Enumeration by ELIspot of interferon gamma (IFNγ)-secreting diabetogenic lymphocytes specific for the proinsulin peptide 15-23 in spleen and pancreatic lymph nodes (pLN). **c)** Enumeration by ELIspot of interferon gamma (IFNγ)-secreting diabetogenic lymphocytes specific for the islet-specific glucose-6-phosphatase catalytic subunit-related protein (IGRP) peptide 206-214 in spleen and pLN. **d)** IGRP-specific CD8+ peripheral blood T cells assessed by staining with the class I MHC tetramer complexed with the NRP mimotope (NRP-V7); **e)** proportion of circulating pathogenic CD8+ T cells specific for IGRP using the class I MHC tetramer complexed with the NRP mimotope (NRP-V7) correlated with the advent of T1D advent in individual mice.

**Figure 4:**
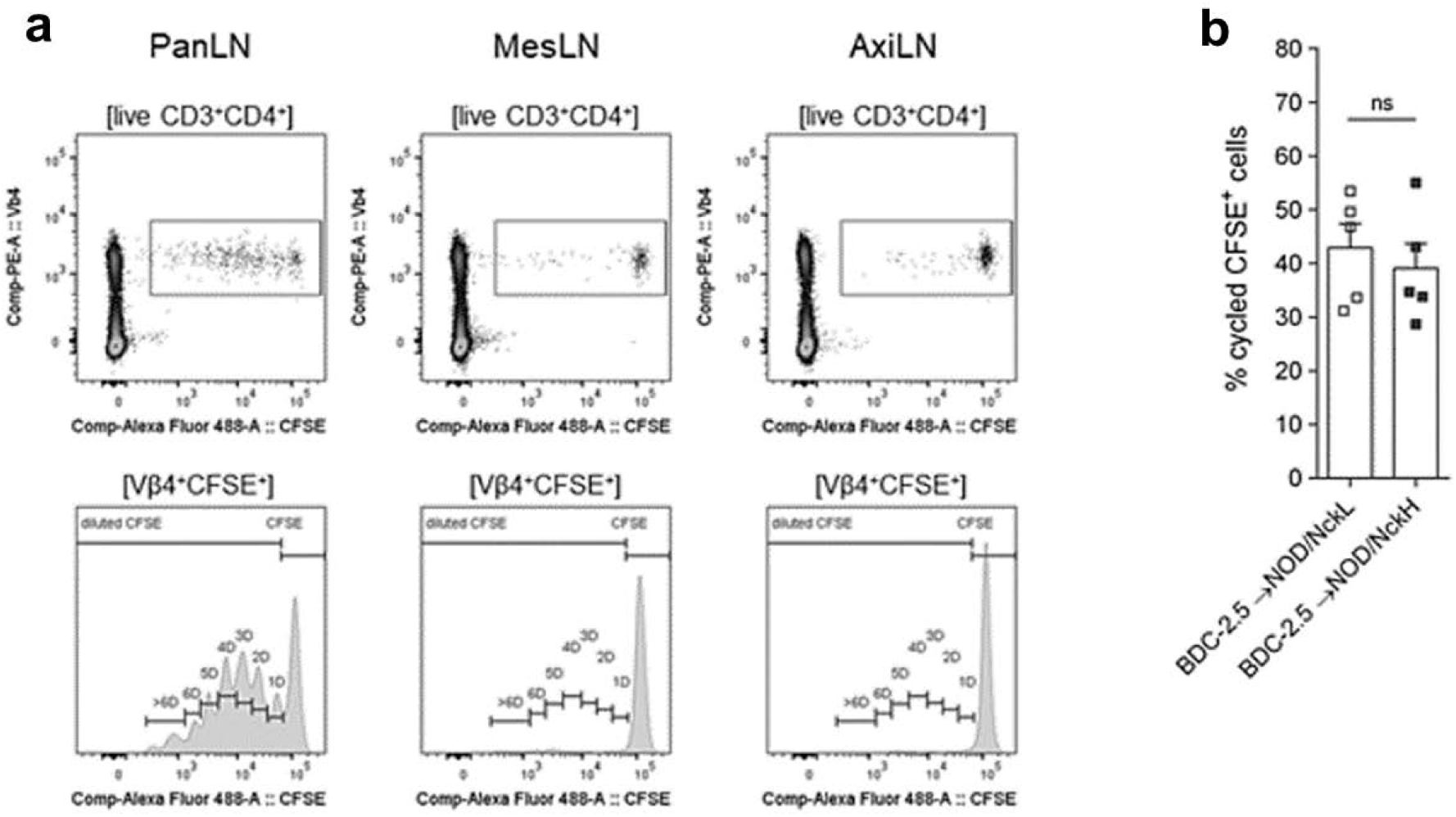
Similar timing of islet antigen presentation in NOD/Nck^H^ and NOD/Nck^L^ mice. **a)** Proliferation of CFSE-labelled transgenic BDC-2.5 CD4+ T cells in pancreatic draining lymph nodes (PanLNs) and in non-draining mesenteric (MesLn) and axillary (AxiLN) lymph nodes of recipient NOD/Nck mice. **b**) Percentage of proliferating cells in 4-week-old NOD mice of the two sublines 96 hours after injection of 10^6^ CFSE-labelled naïve BDC-2.5 CD4+ T cells. Mann-Whitney test; ns, non-significantly different.

Genetic drift is one major factor explaining phenotypic differences within inbred strains; data has been reported in C57BL/6, C3H/HeJ, BALB/c and 129/sV mice (*9–18*). At generation 7 we sequenced the whole genome of four individuals of each subline to search for specific and apparently fixed genetic variation that might contribute to the phenotypic differences in T1D incidence between the NOD/Nck^H^ and NOD/Nck^L^ sublines.

Sequencing data were generated to an average mapped read depth of 31.4X per individual and variants were then called from the alignment against the mm10 reference genome (C57BL/6J; GRCm38). After filtering, a total of 925 putative variants including 745 single nucleotide variants (SNVs), 146 small indels, and 34 structural variants (SVs), were identified. Only two variants were present in insulin-dependent diabetes (*Idd*) genetic regions, a non-coding SNV in *Paqr8* and a non-synonymous missense SNV in *H2-Q4*.

Of the above-mentioned putative variants, 205 were selected for validation, 117 and 88 from the NOD/Nck^H^ and NOD/Nck^L^ sublines, respectively, affecting both coding and non-coding regions. This validation set included 147 SNVs, 21 small indels, and 34 SVs. 118 mutations were validated in 10-15 additional individuals by Ion Torrent sequencing and used for mapping the early T1D onset phenotype observed in the NOD/Nck^H^ subline (Supplementary Table 1). Of the 118 validated variants, 31 were predicted to change coding sense, including 16 non-synonymous missense SNVs. The remaining 15 coding variants were nonsense, stop/start loss, critical splice site mutations, large deletions or duplications, and small indel variants causing a frameshift. Of 14 SVs that validated via Ion Torrent sequencing, only one was found to be unique to the NOD/Nck sublines, as the rest were also present in the NOD/ShiLtJ genome. It was also the only SV to exist in a gene coding region. Transcriptome analysis followed by qPCR validated the 7.4 kb deletion on chromosome 9 overlapping with the coding region of the *Rp9* gene. Interestingly, we found that the validated coding mutations were private to either NOD/Nck^H^ or NOD/Nck^L^ sublines since they were not found in the NOD/ShiLtJ bred since 1988 at the Jackson Laboratory (*14*).

To identify causative mutations, we mapped the autoimmune T1D phenotype as a quantitative trait, age of T1D onset, in a large cohort of F2 hybrid mice made by crossing NOD/Nck^H^ and NOD/Nck^L^ parents and intercrossing their progeny **(Figure 5)**. We combined this traditional breeding approach for quantitative trait locus (QTL) analysis with a validated method for variant genotyping (Ion Torrent sequencing using barcoded libraries) and software tool (Linkage Analyzer) to identify causative mutations present in the germline of mice concurrent with recognition of variant phenotypes (*15*). The program tests the probability of single locus associations with phenotypes using recessive, semi-dominant (additive), and dominant transmission models (*15*). For this particular study, a second mode of analysis was developed in which epistatic effects of mutations at all loci were tested to identify modifications of phenotypic effects by mutations at all other loci.

**Figure 5:**
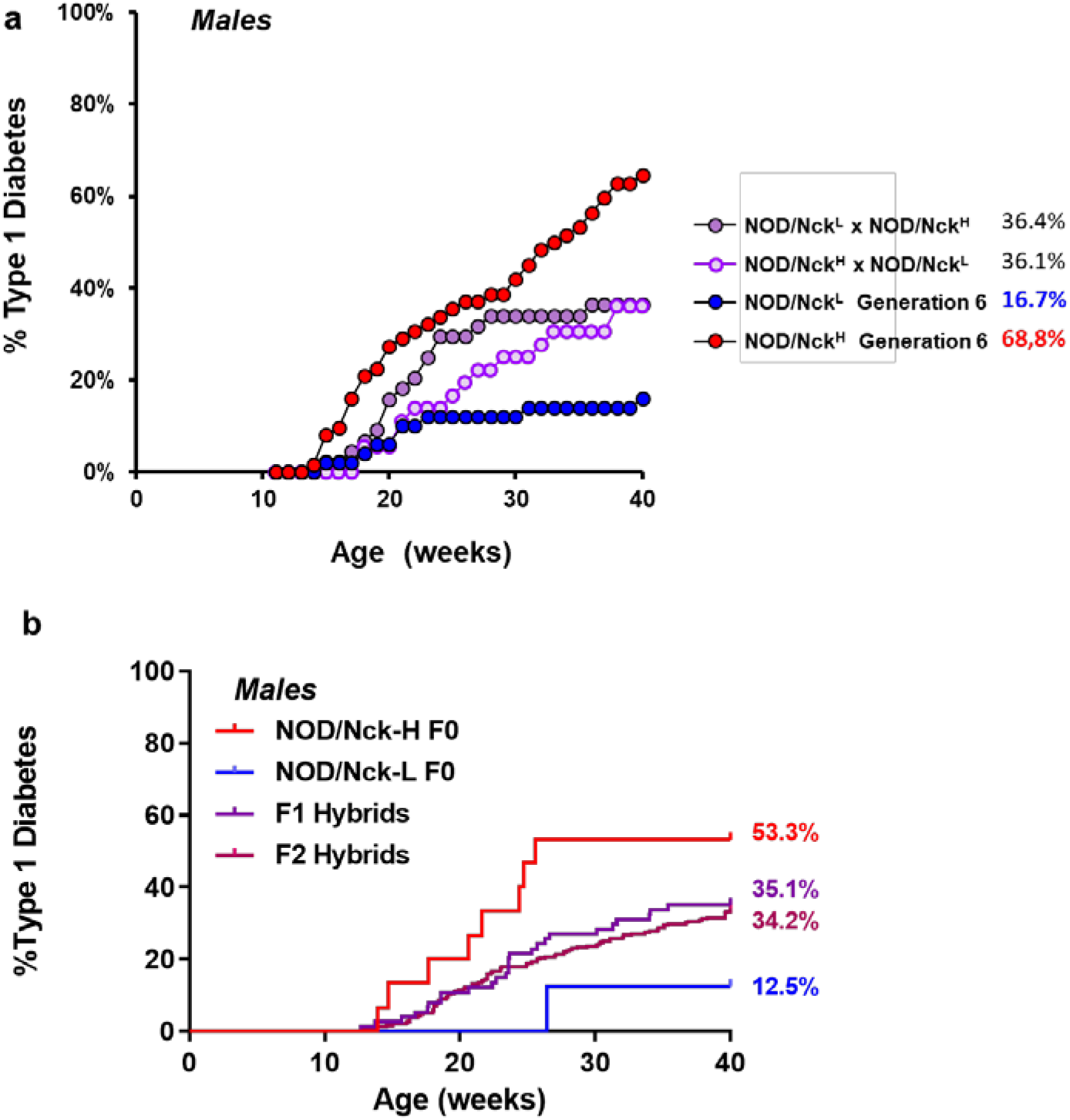
Incidence of T1D in F1 and F2 hybrids of NOD/Nck^L^ and NOD/Nck^H^ mice. **a)** F1 hybrids obtained by crossing NOD/Nck^H^ and NOD/Nck^L^ at generation 6. Incidence of T1D in F1 hybrids is presented with that of the parental lines **b**) Incidence of T1D in F1 hybrids obtained by crossing NOD/Nck^H^ and NOD/Nck^L^ at generation 24, and in F2 hybrids obtained by intercross of F1 hybrids is shown. Data from F2 hybrid mice were used for AMM.

Importantly T1D incidence in F1 hybrids at 40 weeks was intermediate to that of parental F0 NOD/Nck^H^ and NOD/Nck^L^ mice **(Figure 5a,b)**. Also, T1D incidence in F1 hybrids was similar whether the dam was from one or the other subline indicating that there was no subline-specific parental origin effect on the phenotype transmission **(Figure 5a)**. The F2 progeny were generated from intercrosses of F1 NOD/Nck^H^ and NOD/Nck^L^ parents of the 24^th^ generation, and included 294 males and 316 females that showed an incidence and age of onset of T1D similar to that of F1 hybrids at 40 weeks **(Figure 5b)**.

When single locus linkage analysis was performed using Linkage Analyzer the early age of T1D onset in the NOD/Nck^H^ line was mapped to a recessive missense mutation of NOD/Nck^H^ origin in *Dusp10*, encoding the dual specificity phosphatase 10 (DUSP10), also known as mitogen-activated protein kinase (MAPK) phosphatase 5 (MKP5) **(Figure 6)**. Subsequently, an epistatic effect was detected indicating a dominant inhibitory effect of a non-coding polymorphism of NOD/Nck^L^ origin in *progestin and adipoQ receptor family member VIII* (*Paqr8*) on the *Dusp10* variant allele **(Figure 6)**. Importantly, cumulative T1D incidence curves in F2 mice homozygous for both variants strictly reflected the disease incidence curves in parental sublines tested concurrently **(Figure 5b)**.

**Figure 6:**
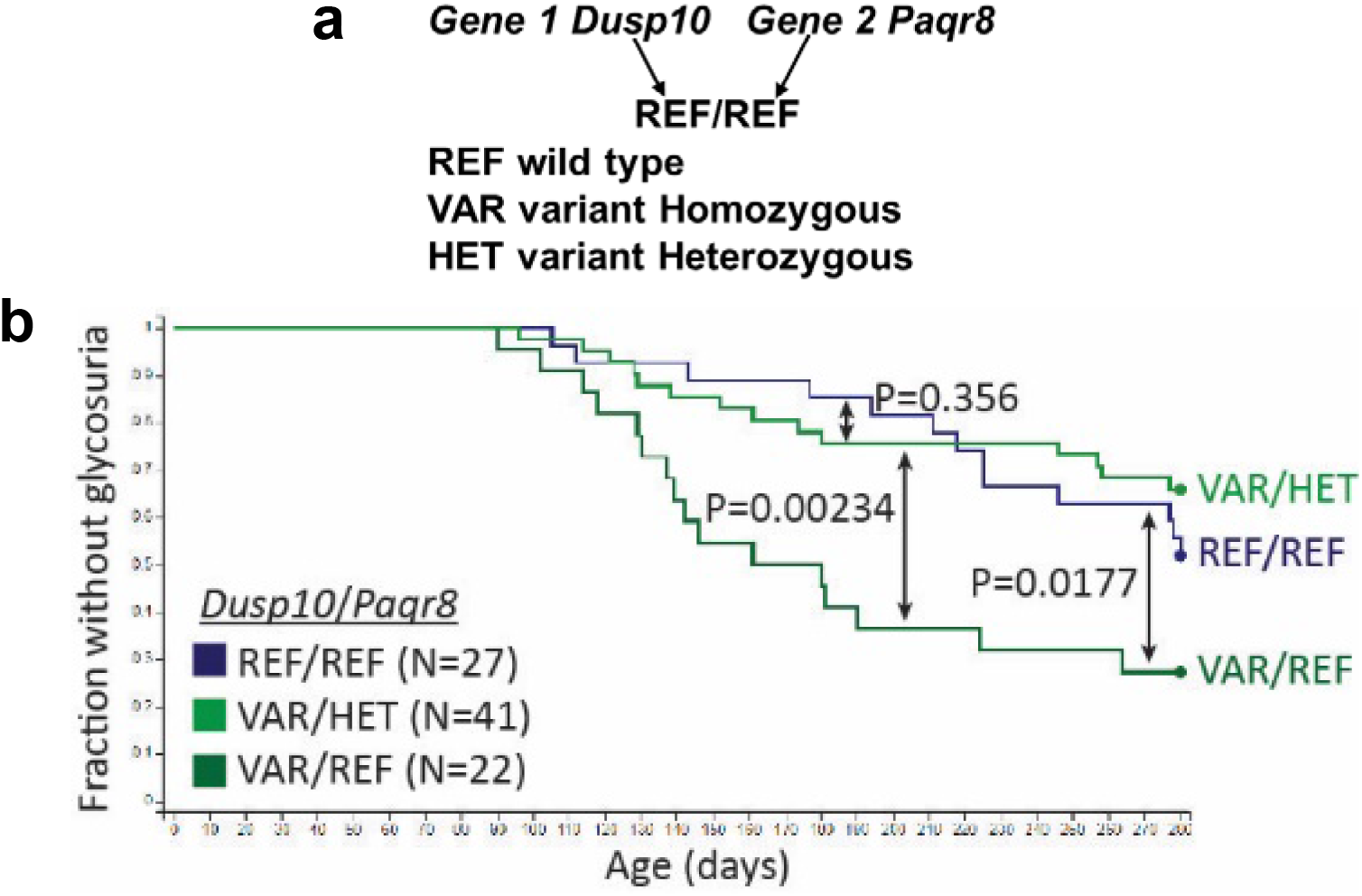
Association of early T1D onset in NOD/Nck^H^ mice wtih a recessive *Dusp10* mutation, and dominant inhibitory effect of a heterozygous *Paqr8* mutation. **a)** For sake of clarity here we describe the nomenclature used in **b)** which designates as indicated by the arrows the allelic forms of *Dusp10* **above** the fraction bar, and the allelic forms of *Paqr8* **below** the fraction bar. REF: homozygous wild-type, VAR: variant homozygous, HET: variant heterozygous. **b)** Kaplan-Meier analysis of glycosuria onset in F2 mice with the indicated *Dusp10* and *Paqr8* genotypes. As compared to wild-type F2 individuals expressing control alleles for *Dusp10* and *Paqr8* (indicated as REF/REF, the blue line), F2 mice expressing the *Dusp10* mutation in the homozygous state (indicated as VAR/REF, the dark green line) showed significant acceleration of T1D development (recessive effect). Moreover, heterozygosity for the *Paqr8* mutation (indicated as VAR/HET, the light green line) abolishes the recessive *Dusp10* effect, rescuing mice from early onset T1D (dominant effect of *Paqr8* mutation).

This *Dusp10* mutation is classified as probably damaging with a Polyphen-2 score of 0.998 (**Figure 7a).** The *Dusp10* nonsynonymous single-nucleotide variant (SNV) changes G to T at base-pair (bp) DUS184,037,056 of chromosome 1, producing a cysteine-to-phenylalanine missense mutation at position 73 in the protein DUSP10 **(Figure 7b)**.

**Figure 7:**
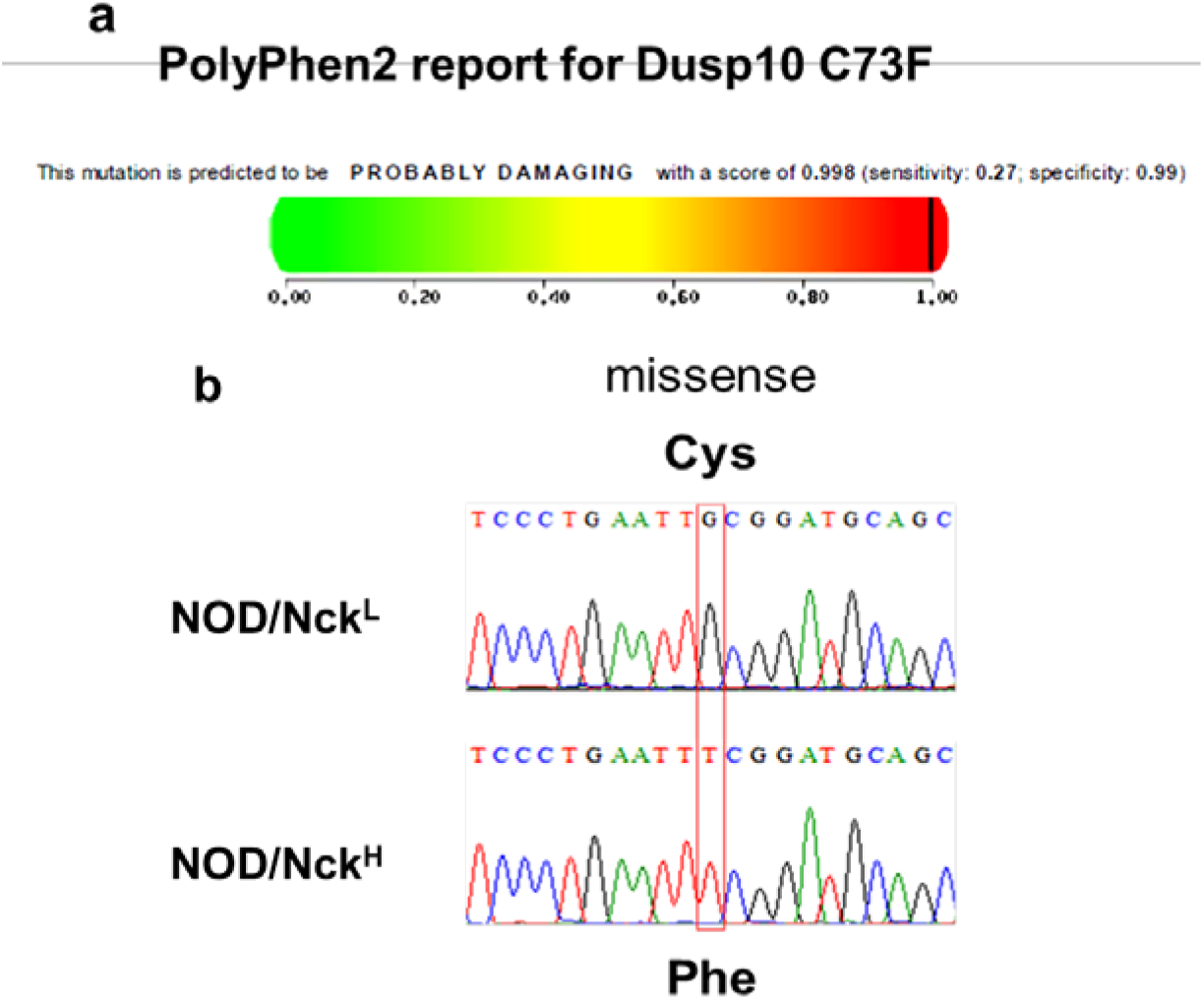
*Dusp10* encodes a Dual Specificity Phosphatase also termed MKP5. **a)** The *Dusp10* mutation we describe is classified as probably damaging with a PolyPhen-2 score of 0.998/1. **b)** The *Dusp10* nonsynonymous single-nucleotide variant (SNV) produces a cysteine-to-phenylalanine missense mutation at position 73 in the protein DUSP10 within an intrinsically disordered N-terminal domain.

MAP kinase phosphatases (MKP), a family of dual-specificity (phospho-Ser/Thr and phospho-Tyr) phosphatases, are essential regulators in immune responses whose function is evolutionarily conserved. These enzymes include three domains. The structures of two of these are known, namely the C-terminal catalytic domain (Residues 330-462 in the *M. musculus* sequence **PF00782.20),** and the central MAPK binding domain (BD, residues 155-280 in the *M. musculus* sequence, also called Rhodanese-like domain PF00581) is involved in recognizing the target MAP kinase (*16, 17*). The variant we describe here is localized in an intrinsically disordered N-terminal domain (from residue 1 to 143, as predicted by Spot-Disorder, alternation of ordered and disordered regions from residue 1 to predicted 140 by PrDos and from residue 1 to 155 by Biomine).

We searched and found highly conserved orthologues of Dusp10 in representatives of all vertebrate classes, including the lamprey *Petromyzon marinus*. The intrinsically disordered N-terminal domain is less conserved than the central and catalytic domains; nevertheless, a stretch of residues including Cys73 (of the *M. musculus* sequence) is completely conserved throughout the vertebrates **(Figure 8)**.

**Figure 8:**
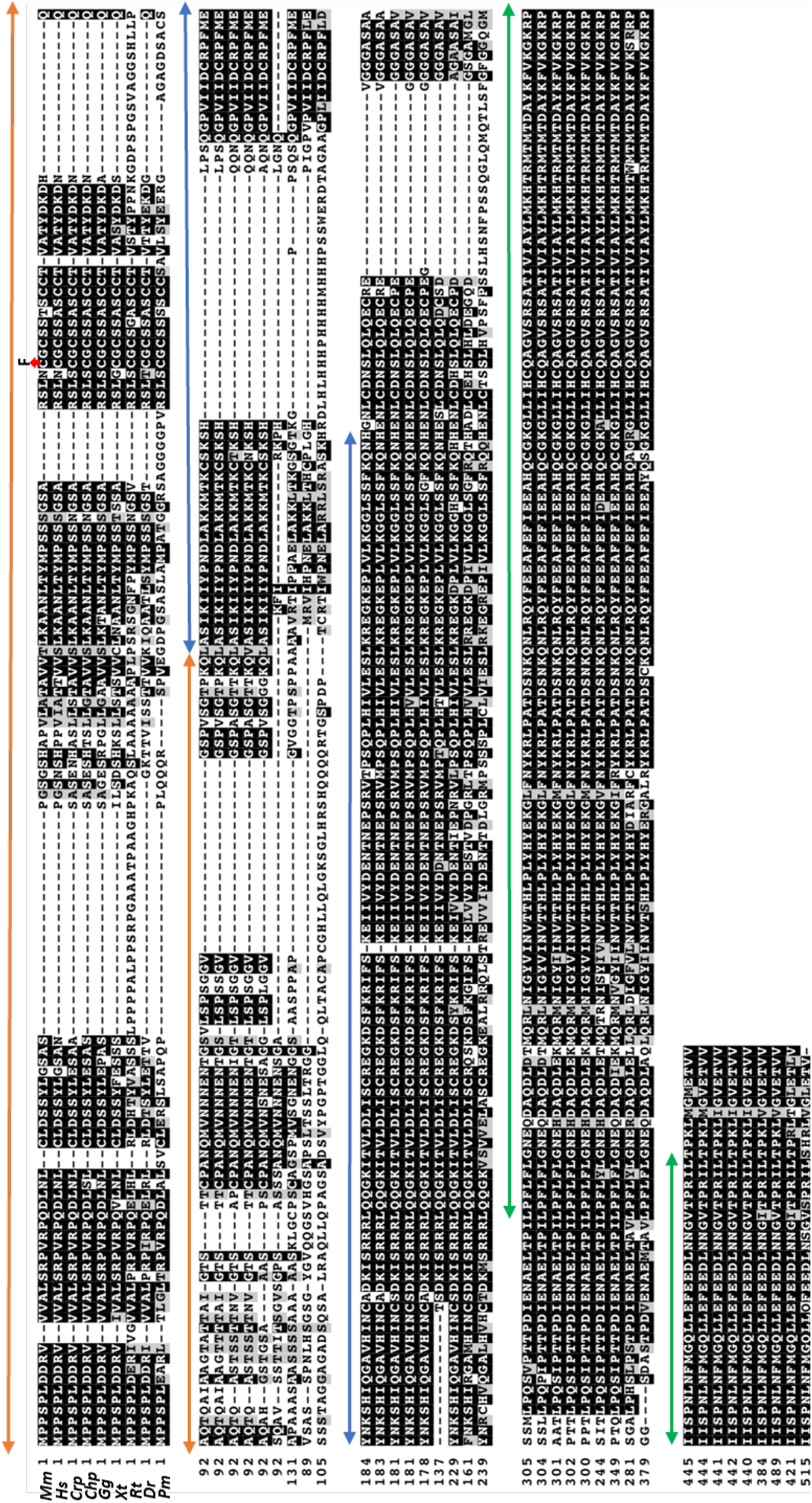
Alignment of putative orthologues of the murine (*Mm*) and human (*Hs*) Dusp10 protein in representative vertebrates. *Crp: Crocodylus porosus*, reptile, crocodile; *Chp*: *Chrysemys_picta_bellii* reptile, turtle (sub-class chelonia); *Gg:Gallus gallus* Aves; *Xt*: *Xenopus tropicali*s, amphibia; *Rt: Rhincodon typus* Chondrichthyes (cartilaginous fishes) Whale shark; *Dr: Danio rerio*, Actinopterygii (sub-class bony fishes); *Pm: Petromyzon marinus* Hyperoartia, Sea Lamprey. ***The red symbol*** indicates the Cys to Phe mutation of the mouse Dusp10 protein discussed in the text. Alignment carried out with MAFFT G-INS-i with default parameters and visualised with BoxShade. ***Double headed arrows*: Orange**: intrinsically disordered sequence, **Blue**: central “Rhodanese-like” domain, **Green**: catalytic domain.

Zhang et al. established C57BL/6 mice homozygous for a robust knockout allele of *Dusp10 (18). Dusp10*-deficient cells showed increased innate immune responses; i.e. production of greatly enhanced levels of pro-inflammatory cytokines by macrophages upon TLR2, TLR3 and TLR4 stimulation (*18*). *Dusp10*-deficient CD4^+^ and CD8^+^ effector T cells produced significantly increased levels of cytokines, which led to much more robust and rapidly fatal immune responses to secondary infection with lymphocytic choriomeningitis virus, contrasting with increased resistance to experimental autoimmune encephalomyelitis (*18*).

To generate a second mutant allele of *Dusp10* on the NOD/Nck^L^ background we used the clustered regularly interspaced short palindromic repeats (CRISPR)/Cas9 technology (*15, 19*). Fertilized one-cell mouse embryos from superovulated NOD/Nck^L^ females mated with NOD/Nck^L^ males were microinjected to target *Dusp10*. We focused our attention on male CR#13 carrying modification *Dusp10*^Y86X^ resulting from the random deletion of a single nucleotide generating a premature STOP codon and potentially leading to a truncated protein with loss of the kinase interaction motif and the phosphatase active site **(Figure 9)**. To reduce the risk of potential off targets effects when generating the CR#13 subline by repeated intercrosses, we first backcrossed the founder to a particular NOD/Nck^L^ subline in which the *Paqr8* polymorphism, which had never become fixed in the homozygous state by inbreeding, was wild-type. This eliminated the dominant inhibitory effect of the *Paqr8* polymorphism on the NOD/Nck^H^ *Dusp10* variant allele **(Figure 10)**. CR#13 mice carrying the homozygous putative null *Dusp10*^Y86X^ mutation showed a T1D incidence similar to that of NOD/Nck^H^ and significantly higher than those homozygous or heterozygous for the wild-type allele whose incidence was similar to that of NOD/Nck^L^.

**Figure 9:**
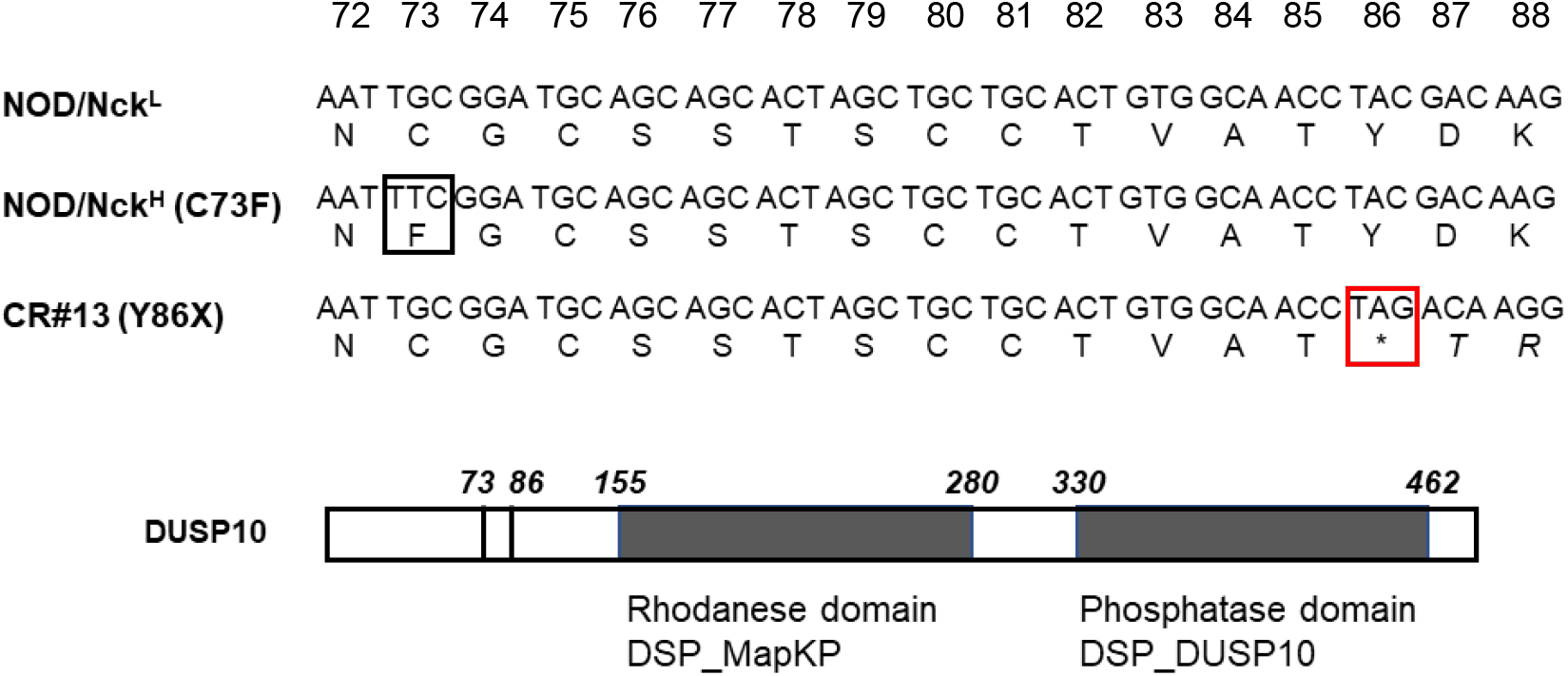
*Dusp10* nucleotide sequence and deduced amino acid sequence. This diagram shows the part of the first *Dusp10* domain containing the nonsynonymous G to T transversion resulting in the cysteine-to-phenylalanine missense mutation at position 73 in the protein DUSP10 (black square) in NOD/Nck^H^ mice. The mutation in CR#13 mice is also shown, here the deletion of a single nucleotide generates a premature STOP codon at position 86 of the protein (red square). Amino acid numbers are shown above.

**Figure 10:**
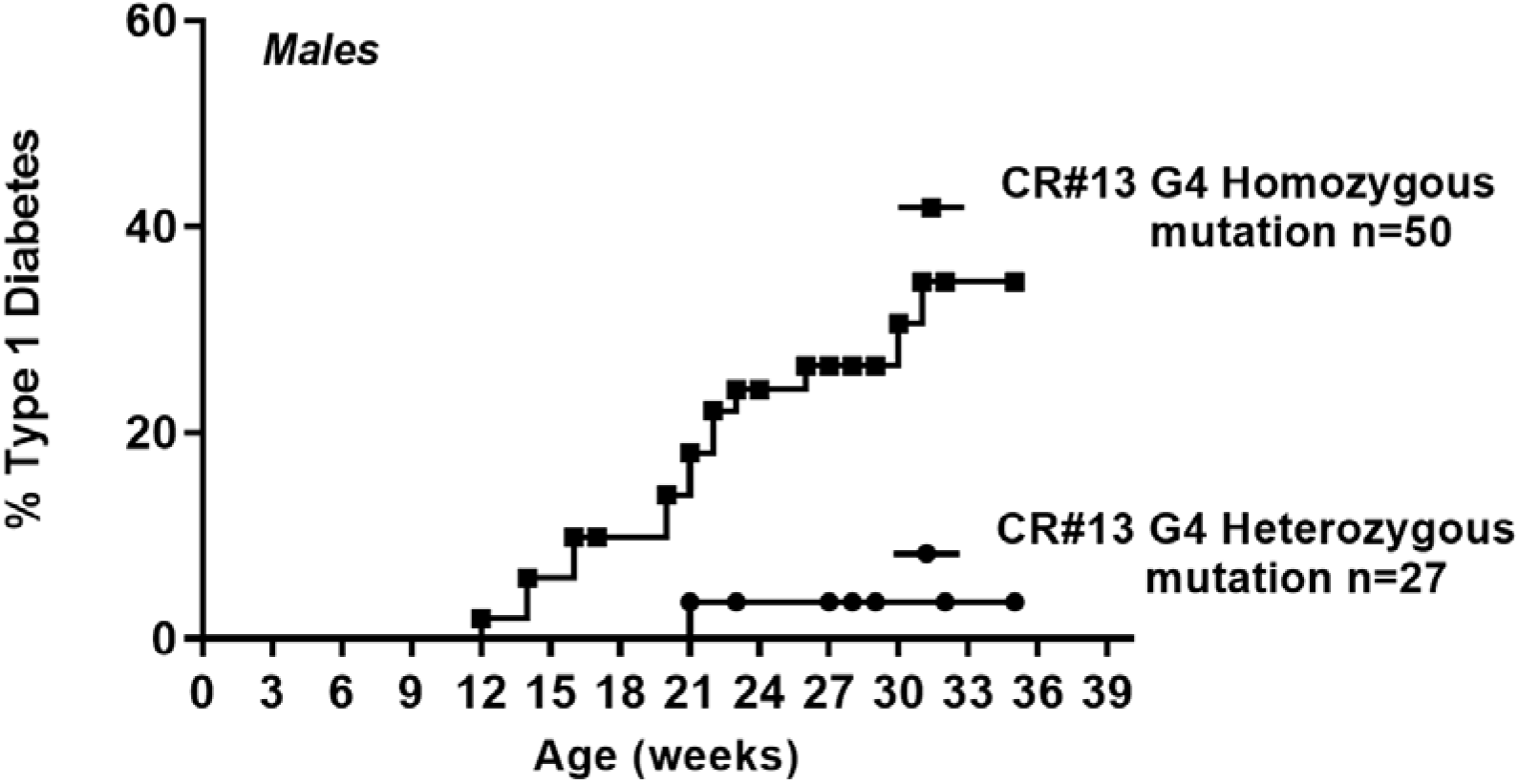
Incidence of T1D in CR#13 subline generated by targeting *Dusp10* with CRISPR/Cas9 in NOD/Nck^L^ mice. Incidence of T1D in male NOD/Nck^H^ and NOD/Nck^L^ mice of generation 34 concurrent to generation 4 of CR#13 was respectively, 53 and 2%.

A fundamental issue was to address whether the difference in T1D incidence driven by the identified *Dusp10* mutation resulted from an impact on the immune system, or on the target of disease (namely the pancreatic islets). To approach this question, we used a strategy described in our laboratory relying on the adoptive transfer of pathogenic T lymphocytes from diabetic animals into newborn syngeneic recipients within the first 3 days of life (*20*). Spleen T lymphocytes from overtly diabetic NOD/Nck^H^ or NOD/NcK^L^ were transferred into the reciprocal newborn recipients. Results showed that NOD/NcK^L^ recipients were less sensitive to T1D transfer than NOD/Nck^H^ recipients arguing for an impact of the *Dusp10* mutation on the islets of Langerhans **(Figure 11)**.

**Figure 11:**
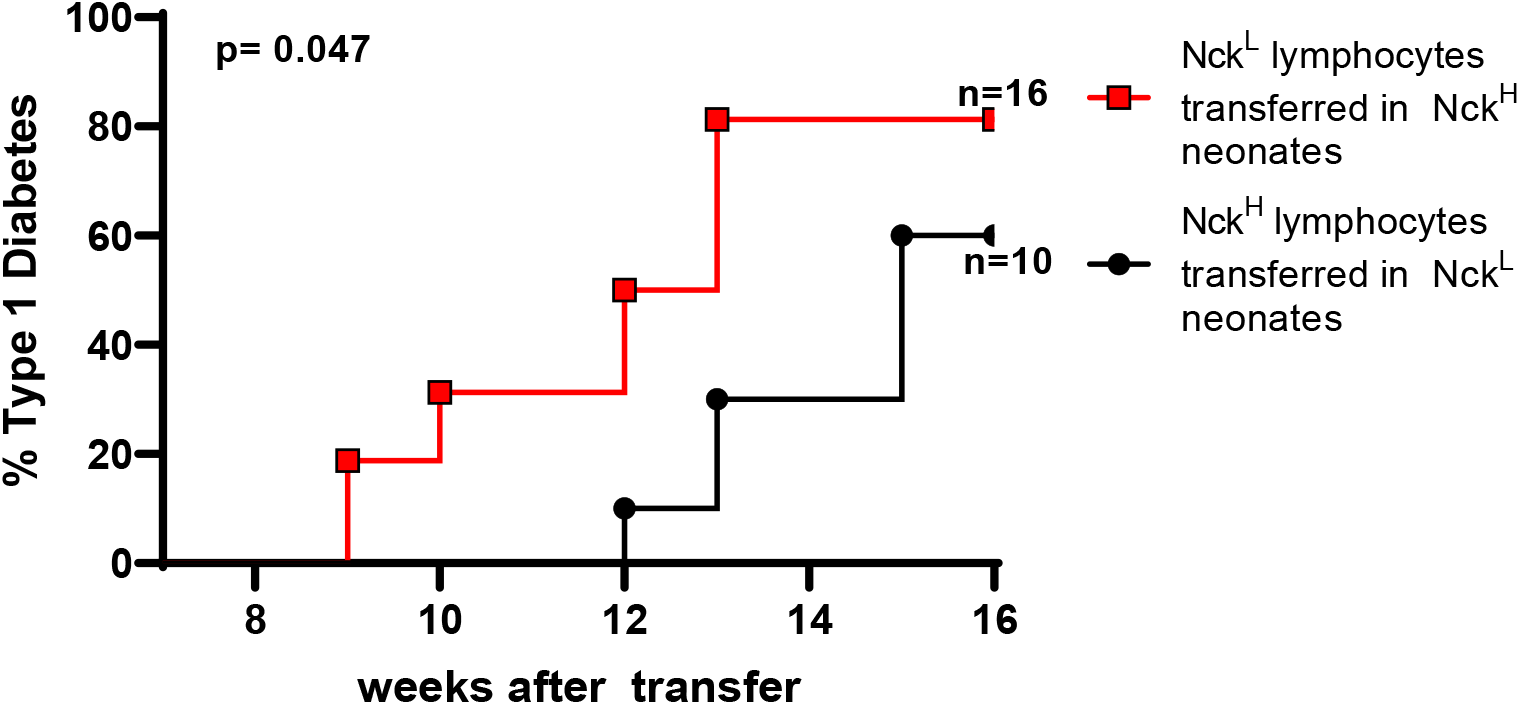
Adoptive transfer of diabetogenic lymphocytes into female NOD/Nck^H^ and NOD/Nck^L^ newborns. Within 24-48 hrs after birth neonates subjected to hypothermic anesthesia (4 min at -20°C) were injected via the periorbital superficial vein under microscopical control with 20 x 10^6^ B cell-depleted splenocytes from diabetic donors. Groups of experimental and control (sham injected) recipients were equally distributed within the litters used. NOD/NcK^L^ recipients were less sensitive to T1D transfer than NOD/Nck^H^ recipients arguing for an impact of the *Dusp10* mutation in the islets of Langerhans.

We performed a comparative transcriptome analysis of several tissues isolated from NOD/Nck^H^ and NOD/NcK^L^ mice **(Figure 12a,b)**. Importantly, pancreatic islets of Langerhans from NOD/Nck^H^ mice isolated at 3 weeks of age, before onset of insulitis, displayed downregulation of 27 genes compared to those in NOD/NcK^L^ islets, including type 1 interferons (*Ifnb*), and various interferon signature genes (*Irgm1, Irgm2, Ifi47, Ifit1, Ifit2, Ifi202b*) and also *Cd274* **(Figure 12b)**. The case of *Cd274*, encoding Programmed Death Ligand 1 (PD-L1) is particularly interesting. The PD-1/PD-L1 axis is an important checkpoint in autoimmune diabetes as assessed by the acceleration of T1D observed in NOD mice upon injection of antibodies to PD-1 (*21*). This result was validated by qPCR on islets isolated from NOD/Nck^H^ and NOD/NcK^L^ **(Figure 12c)**. Immunohistochemical studies in progress show that insulitis, which appears earlier in NOD/Nck^H^ mice, is only observed at early stages in islets expressing low levels of PD-L1. This is in contrast with the high PD-L1 expression observed in insulitis-free islets of NOD/Nck^L^.

**Figure 12:**
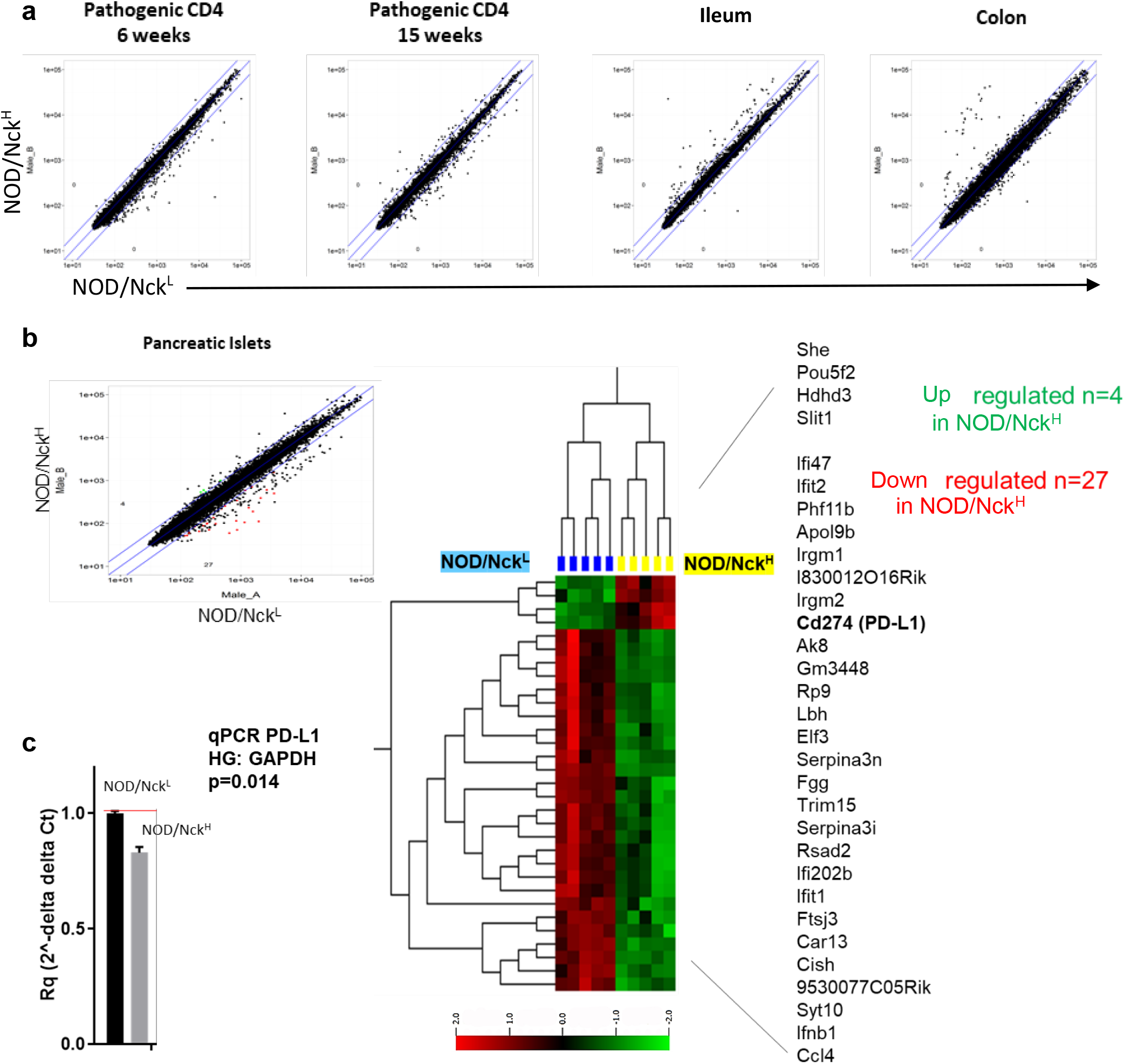
Comparative transcriptome analysis of different tissues from NOD/Nck^H^ and NOD/Nck^L^ mice. **a)** Gene expression in NOD/Nck^H^ versus NOD/Nck^L^ tissues was measured by transcriptome analysis. **b)** Identification of 27 downregulated genes and 4 upregulated genes in purified pancreatic islets from 3-week-old NOD/Nck^H^ mice relative to NOD/Nck^L^ mice. **c)** Increased expression of PD-L1 (CD274) mRNA in NOD/Nck^L^ pancreatic islets was confirmed by qPCR.

Our results have three major implications. First, we identified a single non-synonymous nucleotide change in the DUSP10 encoding gene and provide evidence that it directly affects the phenotype of a complex autoimmune disease. Second, our data illustrate the rapidity of appearance and fixation of *de novo* germline mutations that can alter a complex phenotype such as T1D, either exacerbating it or accelerating it. This suggests that the T1D phenotype has a large genomic footprint. Finally, these findings suggest a new, rapid, and far more precise approach to identifying the missing heritability of T1D and other complex autoimmune diseases, which could be achieved through a forward genetic strategy employing random germline mutagenesis and AMM.

## Materials and Methods

This study was carried out in strict accordance with the recommendations of European Directives (2010/63/UE) and institutional guidelines (INSERM, Faculté Paris Descartes). The protocols were approved by the Ethical Committee of Paris Descartes University and the French Ministry of Education and Research (PROJET N° 2019092612242506 – V3 APAFiS # 25948)

### Mice

NOD/Nck mice were bred and housed under specific pathogen-free conditions at the Hôpital Necker-Enfants Malades animal facility (agreement: C751515). Animals were fed *ad libitum* with an irradiated VRF1 diet (Special Diets Services) with fresh autoclaved water.

### Diabetes monitoring

Mice were monitored for diabetes weekly by testing for glycosuria using colorimetric Diabur-Test 5000 strips (Roche, Boulogne-Billancourt, France). Overt diabetes was confirmed by testing for fasting glycemia >250 mg.dL-1 (Accu-Check; Roche).

### Insulitis scoring

Pancreata were collected, fixed in 4% formaldehyde and paraffin-embedded. Serial 5-μm sections cut at 60 μm intervals were hematoxylin and eosin stained and at least 80 islets per pancreas and/or 200 islets per group of mice were scored. Mononuclear cell infiltration of the islets was graded as follows: 0, no insulitis (intact islet); I, periinsulitis (infiltration remaining confined to the periphery of the islet); II, invasive insulitis (infiltration <50% of the islet surface); III, destructive insulitis (infiltration totally disrupting the morphology of the islet).

### IFNγ producing autoreactive CD8+ T cells detection

For IFNγ ELISpot assay, 96-well PVDF plates (Millipore) and an anti-IFNγ capture and detection antibody pair (U-CyTech Biosciences) were used. Cells were cultured in the presence of IL-2 at 2.5.105 per well in the case of splenocytes or, in the case of lymph node cells, at 0.3.105 per well together with 2.0.105 irradiated (35 Gy, X-Ray source) splenocytes. Peptides used were InsB15-23 and IGRP206-214 (7 μM) and as a positive control was used CD3 antibody (clone 2C11; 1 mg.mL-1). After a 20-hour culture, IFNγ was detected using biotinylated anti-IFNγ capture antibody (U-CyTech Biosicences), alkaline phosphatase conjugated ExtrAvidin (Sigma-Aldrich) and 5-bromo-4-chloro-3-indolyl phosphate/nitro blue tetrazolium (Sigma-Aldrich). Air-dried plates were read, and spots counted using an AID reader (Autoimmun Diagnostika). Data were means of triplicate wells minus the average number of spots in negative control wells (cells cultured without peptides) and were expressed as spot-forming cells per 10^6^ cells.

### Tetramer staining

Peripheral blood collected from the retro-orbital sinus using heparinized capillary tubes was incubated with H-2Kd tetramers bearing the peptide NRP-V7 or an irrelevant peptide (TUM) for 1 hour on ice. Tetramers were provided by P. Santamaria (Calgary University, Canada). After a washing step, surface staining was performed at 4°C in staining buffer (PBS 1X-FCS 1%-Sodium azide 0.1%) using anti-CD8-APC (clone 53-6.7) and anti-CD45R-PerCP (clone RA3-6B2), (both purchased from BD Biosciences). Red blood cells were lysed at room temperature using FACS Lysing Solution (BD Biosciences) before cell suspensions were washed prior to acquisition. Data were acquired on a FACSCanto II flow cytometer (BD Biosciences) and analyzed using FlowJo software (Tree Star). NRP-V7 tetramer-positive cells were expressed as percentage of B220-CD8+ cells minus the percentage of TUM tetramer positive-cells.

### *In vivo* tracking of naïve CD4+ T cell activation

Naïve CD4+ T cells were isolated from spleens and lymph nodes of BDC-2.5 transgenic NOD mice using the Mouse CD4 Naïve T Cell Isolation Kit (BioLegend). Naïve T cells were further selected positively by treatment with anti-CD62L biotinylated antibodies (BD Pharmingen) followed by anti-biotin magnetic beads and sorted using MACS technology (Miltenyi Biotec). After CFSE-labelling (5 μM), 106 cells were injected intravenously into 25-day-old recipients. On day 3, pancreatic, mesenteric and axillary lymph nodes were collected and cells were stained for flow cytometry analysis.

### Adoptive transfer of diabetes into neonates

Within 24-48 hr after birth neonates subjected to hypothermic anesthesia (4min at −20°C) were injected via the periorbital superficial vein under microscopical control with 0.05ml cell suspension at the appropriate concentration (20 x 10^6^ B cell-depleted splenocytes from diabetic donors). Groups of experimental and control recipients (NOD/Nck^H^ and NOD/Nck^L^ newborns sham injected with saline) were equally distributed within the litters used. Here, to ask if the mutation in recipient’s islet of Langerhans cells can impact T1D, NOD/Nck^H^ and NOD/Nck^L^ neonates were injected with 20 x 10^6^ B cell-depleted lymphocytes from overtly diabetic NOD/Nck^L^ and NOD/Nck^H^ mice, respectively.

### Isolation of pancreatic islet cells

Pancreata were perfused with a solution of collagenase P (0.6 mg.mL-1; Roche), dissected free from surrounding tissues and then digested at 37°C for 13 minutes. After extensive washes, islets were purified by handipicking before being dispersed using a non-enzymatic Cell Dissociation Solution (Sigma-Aldrich). Single-cell suspensions were then prepared for flow cytometry as described below.

### Flow cytometry

Single-cell suspensions were prepared from lymphoid organs by and from different tissues. Prior to staining, cells were incubated with a Fixable Viability Dye (eBioscience) and then with 2.4G2 antibody for FcγRII/III blocking (BD Pharmingen). Surface staining was performed at 4°C in staining buffer (PBS 1X-FCS 2%-EDTA 5mM). Intracellular staining of cytokines was performed using Fixation Buffer (BioLegend) and Intracellular Staining Perm Wash Buffer (BioLegend). Transcription factors were stained using the Foxp3 TF Staining Buffer Set (eBioscience). See the list of antibodies at the end of the Materials and Methods section. Data were acquired on a FACSFortessa flow cytometer (BD Biosciences) and analyzed using FlowJo version 10 software (Tree Star).

### Whole-genome sequencing and variant identification

High molecular weight genomic DNA was prepared from the liver of 4 animals of each NOD/Nck subline. Whole-genome sequencing was performed using an Illumina HiSeq 2500 (Illumina, San Diego, USA). Sequencing data were mapped using BWA (v 0.7.10) to align against the mm10 reference genome (C57BL/6J; GRCm38). Using the Genome Analysis Toolkit (GATK) and SAMtools we found an average of 7.3 million single nucleotide variants (SNV) and small insertions-deletions (indels) differentiating the NOD/Nck and the C57BL/6J strains using single-sample variant calling. For purposes of filtering, joint variant calling using the four samples of each subline was performed. Using this jointly called data, variants from each subline were compared. They were then filtered for those variants unique to each subline and heterozygous calls excluded (allelic ratio > 0.875). A quality score of 80 was used as a filtering threshold for jointly called variants. Additionally, 34 SVs differentiating the NOD/Nck and C57BL/6J strains were identified using speedSeq SV (v 0.1.0). In addition to the SVs, 168 SNVs and small indels were chosen for validation via Ion Torrent sequencing to give genomic linkage using 20Mb as a cutoff. In parallel, alignment of sequencing data to mouse reference genome (GRCm38, mm10), variant calling, and automated filtering were performed by collaborators (M. Dumas and B. Jost, Plateforme biopuces et séquençage, IGBMC, Strasbourg, France) with particular attention to variants in non-coding regions. Manual checking was performed using an IGV2.3 viewer (Broad Institute). A total of 118 variants were validated in 10-15 additional mice and used for AMM. 77.3% of the mouse genome was in linkage with at least one of the 118 sequence variants, using a distance cutoff criterion of 20 Mb.

### Automated meiotic mapping (AMM)

A panel of 118 validated variant loci distinguishing NOD/Nck^H^ and Nod/Nck^L^ sublines was used to map the high diabetes incidence phenotype in F2 mice. The F2 animals generated by two-generation intercrosses were monitored for overt diabetes for 40 weeks to determine the age of diabetes onset. Genomic DNA was extracted from tail snips and used as a template for multiplexed targeted amplification of the 118 loci using custom Ampliseq primers. Amplification products were barcoded to correspond to individual mice prior to sequencing using an Ion PGM (Life Technologies). Following phenotypic screening, AMM using recessive, additive, and dominant models of inheritance was performed for each variant using the Linkage Analyzer program as previously described (*15*). Kaplan-Meier plots of phenotypic data and Manhattan plots of linkage data were generated using the Linkage Explorer program (*15*). The *P* values of association between genotype and phenotype were calculated based on Kaplan-Meier analysis of time of onset of T1D, as related to zygosity for each of the mutations using a likelihood ratio test from a generalized linear model or generalized linear mixed effect model and Bonferroni correction applied.

### Generation of genetically modified NOD/Nck mice using CRISPR/Cas9

NOD females were superovulated by administration of PMSG at day 0 (5U) and hGC at day 2 with a 44 hour-interval (5U), before being mated with NOD males. Because NOD females were found to give more eggs at 8 weeks of age than at 4 weeks of age (data from Fabrice Valette), 8-10-week-old mice were used as zygote donors. Zygote collection, fertilized egg microinjection, and transfer in the oviduct of pseudogestant foster mothers were performed by collaborators (Plateforme de Transgénèse, F. Langa-Vives, Institut Pasteur, Paris). The design of the single-guide RNA for the knock-in and knock-out experiments, and the selection of the HDR donor template for the knock-in experiment, were performed in collaboration with the laboratory of B. Beutler (UT Southwestern, Dallas, USA). All the reagents were from Integrated DNA Technologies (IDT): crRNA (IDT Alt-R CRISPR-Cas9 crRNA), tracrRNA (IDT Alt-R CRISPR-Cas9 tracrRNA), HDR donor template (IDT Ultramer DNA oligo), and Cas9 (IDT Alt-R S.p. HiFi Cas9 Nuclease 3NLS). For genotyping of pups, DNA was extracted from tail snips using Genomic DNA purification kit (Qiagen) and proteinase K (Roche). PCR was performed by using the forward *Xf* and reverse *Xr* primers and Accuprime *Pfx* DNA polymerase (Invitrogen) and the PCR products, after purification using the QIAquick PCR purification kit (Qiagen), were analyzed by sequencing using *Xf* primer (Plateforme de séquençage, Eurofins).

### Transcriptome Gene expression profiling

Agilent SurePrint G3 Mouse Gene Expression 8×60K Microarrays (Agilent Technologies, Santa Clara, CA, USA) were used for microarray experiments on purified CD4+CD25-CD62L-spleen lymphocytes, colon, ileon and islets of Langerhans from NOD/Nck^H^ and NOD/Nck^L^ mice. Five replicas were prepared for the various cell/tissue preparations. Cell and islet pellets and tissues were lysed using SuperAmpTMlysis buffer (Miltenyi Biotech, Waldbronn, Germany) and stored at −80°C until all samples were collected. Preparation of RNA, amplification of RNA, sample hydridization (Agilent whole mouse genome oligo microarrays), scanning and data acquisition were performed by Miltenyi Biotech. RNA quality and purity were assessed on the Agilent Bioanalyzer 2100 (Agilent Technologies). Microarray probe fluorescence signals produced by the Agilent Feature Extraction (AFE) image analysis software were converted to expression values using Bioconductor *Agi4×44PreProcess* package with a custom annotation package.

### Antibodies for staining

Biotin or fluorochrome-labeled antibodies to murine CD11c (HL3), B220 (RA3-6B2), CD8 (53-6.7), CD103 (2E7), CD80 (19-10A1), CD86 (GL1), CD40 (3/23), I-Ak (10-3.6; crossing with the MHC class II from NOD mice, I-A^g7^), CD11b (M1/70), ICAM-1 (3E2), OX40L (RM134L), CCR7 (4B12), CCR9 (9B1), TCR (H57-597), CD4 (GK1.5), CD19 (1D3), CD25 (PC61), DX5 and streptavidin-pacific blue were obtained from PharMingen BD. Antibodies to PD-L1 (MIH5), GITRL (eBioYGL386), CD73 (eBioTY/11.8), CD39 (24DMS1) were purchased from eBisocience.

### Statistical analyses

All statistics were performed using GraphPad Prism version 6 software. Cumulative actuarial diabetes incidence was calculated according to the Kaplan-Meier method. Incidence curves were compared using the log-rank (Mantel-Cox) test. Comparison between means was performed using the non-parametric Mann-Whitney test for other experiments, unless otherwise specified in the figure legend. Data are presented as means ± S.E.M. A *P* value <0.05 was considered statistically significant; \**P* <0.05, \*\**P* <0.01, \*\*\**P*<0.001, \*\*\*\**P*<0.0001.

## Acknowledgements

This work was supported by National Institutes of Health grants R01 AI125581 and U19 AI100627 (to B.B.), and by an Advanced grant from the European Research Council (ERC, Hygiene N°: 250290) (to J-F.B.), and by Institut National de la Santé et de la Recherche Scientifique (INSERM), Fondation Day Solvay, and the Lyda Hill Foundation. The authors are indebted to Dr. Sylvaine You, Dr. Alicia Pérez-Arroyo, Mrs. Laurène Magne, Mme. Céline Keime, and Mr. Bernard Jost for their support of this work.

## Author contributions

C.S., J-F.B., B.B., L.C. designed research; A-P.F., S.C., C.M., F.V., C.P., S.Lemoine, P.H., F.L-V., L.S. performed research; P.S., S.Lyon, C.H.B., T.W., D.X. contributed new reagents/analytic tools; A-P.F., S.C., M.D., S.H., S.Lemoine, C.H.B., C.S., L.C. analyzed data; A-P.F., S.C., C.S., J-F.B., B.B., L.C. wrote the paper; E.M.Y.M., J-F.B., B.B., L.C. revised the paper.

## References

1. J. F. Bach, Insulin-dependent diabetes mellitus as an autoimmune disease. Endocrine Rev. 15, 516–542 (1994).

2. W. M. Ridgway et al., Gene-gene interactions in the NOD mouse model of type 1 diabetes. Advances in immunology 100, 151–175 (2008).

3. J. F. Bach, The effect of infections on susceptibility to autoimmune and allergic diseases. N Engl J Med 347, 911–920 (2002).

4. P. Concannon et al., Genome-wide scan for linkage to type 1 diabetes in 2,496 multiplex families from the Type 1 Diabetes Genetics Consortium. Diabetes 58, 1018–1022 (2009).

5. S. Makino et al., Breeding of a non-obese, diabetic strain of mice. Exp. Anim. 29, 1–13 (1980).

6. P. Simecek et al., Genetic Analysis of Substrain Divergence in Non-Obese Diabetic (NOD) Mice. G3 (Bethesda, Md.) 5, 771–775 (2015).

7. L. Yurkovetskiy et al., Gender bias in autoimmunity is influenced by microbiota. Immunity 39, 400–412 (2013).

8. J. D. Trudeau et al., Prediction of spontaneous autoimmune diabetes in NOD mice by quantification of autoreactive T cells in peripheral blood. The Journal of clinical investigation 111, 217–223 (2003).

9. V. Kumar et al., C57BL/6N mutation in cytoplasmic FMRP interacting protein 2 regulates cocaine response. Science (New York, N.Y.) 342, 1508–1512 (2013).

10. A. Poltorak et al., Defective LPS signaling in C3H/HeJ and C57BL/10ScCr mice: mutations in Tlr4 gene. Science (New York, N.Y.) 282, 2085–2088 (1998).

11. M. M. Simon et al., A comparative phenotypic and genomic analysis of C57BL/6J and C57BL/6N mouse strains. Genome biology 14, R82 (2013).

12. E. Zurita et al., Genetic polymorphisms among C57BL/6 mouse inbred strains. Transgenic research 20, 481–489 (2011).

13. L. J. Sittig et al., Phenotypic instability between the near isogenic substrains BALB/cJ and BALB/cByJ. Mammalian genome: official journal of the International Mammalian Genome Society 25, 564–572 (2014).

14. T. M. Keane et al., Mouse genomic variation and its effect on phenotypes and gene regulation. Nature 477, 289–294 (2011).

15. T. Wang et al., Real-time resolution of point mutations that cause phenovariance in mice. Proc Natl Acad Sci U S A 112, E440–449 (2015).

16. X. Tao, L. Tong, Crystal structure of the MAP kinase binding domain and the catalytic domain of human MKP5. Protein Sci 16, 880–886 (2007).

17. Y. Y. Zhang, J. W. Wu, Z. X. Wang, A distinct interaction mode revealed by the crystal structure of the kinase p38α with the MAPK binding domain of the phosphatase MKP5. Science signaling 4, ra88 (2011).

18. Y. Zhang et al., Regulation of innate and adaptive immune responses by MAP kinase phosphatase 5. Nature 430, 793–797 (2004).

19. L. Cong et al., Multiplex genome engineering using CRISPR/Cas systems. Science (New York, N.Y.) 339, 819–823 (2013).

20. A. Bendelac, C. Carnaud, C. Boitard, J. F. Bach, Syngeneic transfer of autoimmune diabetes from diabetic NOD mice to healthy neonates. Requirement for both L3T4+ and Lyt-2+ T cells. The Journal of experimental medicine 166, 823–832 (1987).

21. M. J. Ansari et al., The programmed death-1 (PD-1) pathway regulates autoimmune diabetes in nonobese diabetic (NOD) mice. The Journal of experimental medicine 198, 63–69 (2003).

